# Spatiotemporal growth pattern during plant nutation implies fast dynamics for cell wall mechanics and chemistry: a multiscale study in *Averrhoa carambola*

**DOI:** 10.1101/2022.02.22.481493

**Authors:** Mathieu Rivière, Alexis Peaucelle, Julien Derr, Stéphane Douady

## Abstract

Nutation is the most striking and ubiquitous example of the rhythmic nature of plant development. Although there is a consensus that this wide oscillatory motion is driven by growth, its internal mechanisms have not been fully elucidated yet. In this work, we study the specific case of nutation in compound leaves in the archetypal *Averrhoa carambola* plant. We quantify the macroscopic growth kinematics with time lapse imaging, image analysis and kinematics modeling. We further characterize the mechanical and chemical properties of the cell wall with atomic force microscopy and immunolabelling. Our data first reveal that the differential growth driving nutation is localized and peaks where the average growth drops. We then show this specific spatiotemporal growth profile is compatible with local contraction events. At the cell wall level, differential growth is further colocalized with an asymmetry of the cell wall elastic modulus, and with an asymmetric distribution of homogalacturonans (HG). Our results not only back up the hypothesis of HG being involved in plant growth, but also build up on it by suggesting a dynamic nature for this process.

**Significance Statement:** Nutation is an oscillatory motion displayed by many organs of growing plants. Most works on nutation focus on its relation to external stimuli attempting to explain its origins. By contrast, its internal physiological mechanisms remain to be fully explored. Here we propose an experimental and multiscale characterization of undisturbed nutation. We determine the macroscopic growth profile and show it is compatible with cell expansion but also local contractions in the tissues. At the microscopic level, we reveal that both the rigidity and composition of the cell wall are asymmetrically distributed where nutation occurs. The combination of results on both scales brings contributions to the understanding of interplay between global movement, local growth, cell wall mechanics and cell wall biochemistry.

---

**P**lants move. This understated truth has been recently reemphasized by the study of spectacular ultra rapid motions (1). For example, both the snapping of the Venus flytrap (2, 3) or the catapulting of fern spores (4) require high speed cameras to be recorded. At the opposite side of the timescales spectrum, plants moves through their growth. These slow motions necessitate time-lapse imaging. After Darwin (5), they started to be historically investigated with the development of photography (6). But we are still evidencing nowadays a variety of exciting new motions (7–9). They can either be nastic motions, or tropisms, depending on whether the direction of the motion is imposed by factors internal or external to the plant respectively. The trigger can be autonomic or paratonic depending on whether the leading cause of the motion is within the plant or not. They can finally be reversible or linked to irreversible growth. These three dichotomies define the traditional classification of slow plant motion (7). Within this framework, the status of one remarkable movement called nutation is still undecided (7, 10–12)

Nutation is the phenomenon that causes the orientation of the long axis of an elongated growing plant to vary over time in a pseudo-periodical way. It was already observed for climbing plants by British botanists of the 17th century (13) and began to be studied by Hugo von Mohl and Ludwig Palm in the first part of the 19th century (14). To the best of our knowledge, the term “nutation” was first mentioned by Charles Bonnet (15) although he acknowledges that this term had been named before him, by physicists who knew the phenomenon. They probably saw this motion as a botanical analog to the astronomical nutation^*^. Darwin introduced the idea that nutation had an endogenous origin and many plant motions were actually modified nutations (5). The very origin of nutation was a source of debate at the time nonetheless (14), and it remains so up to this date (12, 17, 18). Part of the community backs up Darwin’s idea of an internal oscillator (19, 20). Others ascribe this oscillating behaviour to inertial overshooting of the plant occurring during its straightening process (21–24). Finally, the compromise solution calling for a combination of these two hypotheses attracts more and more attention (10, 20, 25–27). The one thing making consensus is that nutation is obviously linked to growth.

Plant growth results from a subtle balance between the strong internal osmotic pressure and the resisting rheology of the cell wall (28). The viscoplastic framework formalised by Lockhart (29) received good experimental support at the single cell level (30–32). Still, some shortcomings need to be addressed (33), and the origin of the cell wall-loosening mechanism is still unclear (34–37). The cell wall is considered here to be an inactive gel but it was demonstrated that elements of the cell wall, the homogalacturonans (HG) can transform chemical modification into mechanical expansion through cell control enzymatic demethylesterification. This cell wall expansion was demonstrated to be sufficient to recapitulate *Arabidopsis* cotyledon epidermal morphogenesis (38). In this view, HG demethylesterification would be necessary and sufficient for tissue cell growth and even to correlate with changes in cell wall elasticity. The precise role of elasticity that was added to Lockhart model later on by Ortega (39) is then subject to debate (38, 40). Finally, the multi-cellular aspect of the biophysics of growth remains to be formalized and understood (41).

Here we want to investigate in detail the phenomenon of nutation with a double goal in mind. By carefully quantifying the motion of nutation, we will gain knowledge on the nature of this puzzling mechanism; and, because motions are a direct manifestation of differential growth, we will use the nutation motion as a particular proxy to get insight on the very nature of plant growth.

In this article, we present a multi-scale study of nutation in the plant *Averrhoa carambola*, a plant known for exhibiting ample growth motions (7, 8, 42). Its compound leaf first unfurls, creating a hook, that show some circadian modulation and will only disappear at the end of growth (for anatomical terms, please refer to Fig. S1). Along this hook, the folded leaflets reverse their position ab-axially, then unfold, and when coming out of the hook start to reach an horizontal position with a circadian oscillation. On top of these motions the main rachis oscillates from right to left in a nutation motion. Focusing on this nutation motion, we start to characterize it at the plant level. We characterize its dynamics, and emphasize its spatial localization. We continue by focusing on this specific zone which happens to coincide with the distal end of growth zone. Our measurements allow to characterize the growth law of nutation and highlight a relationship between differential growth and average growth. Finally, we dive into the microscopic properties of the plant cell wall by performing atomic force microscopy (AFM) measurements as well as immunohistochemistry. We observed that the cyclic asymmetric growth correlates with decrease cell wall elasticity, accumulation in both highly methylesterified and highly demethylesterified HG but not for sparsely demthylated HG and other cell wall component.

## Results

### Characterizing nutation

As they grow, *Avherroa carambola* compound leaves exhibit pronounced growth motions. Putting aside the leaflets, the motion of the rachis can be broken down into two different motions, depending on their plane of occurrence. The unfurling motion of the rachis of *Avherroa carambola* mostly takes place in a principal plane (7). The rachis unfolds steadily while propagating a hook shape (8). This hook shape is visible in Fig. 1A. This motion is also accompanied by out-of-plane curvature variations. The rachis bends and unbends in a pseudo-periodical way, as if it were oscillating around a rectilinear state. The oscillations can already be seen in Fig. 1A. In Fig. 1B we see the same motion from the top, and on a slightly longer time range. The period of oscillation varies greatly between 1.5 and 4 hours, typically between 2 and 3 hours, while the typical amplitude is of the order of 25 degrees. **Supporting movie 1** shows a time-lapse movie of a typical nutation motion, seen from both sides. To properly describe the nutation motion, we define the arc length *s* going from the base towards the apex^†^, the local angle *ϕ* with respect to the average direction of the rachis and the curvature *K*_⊥_ (see Fig. 1C).

**Fig. 1.**
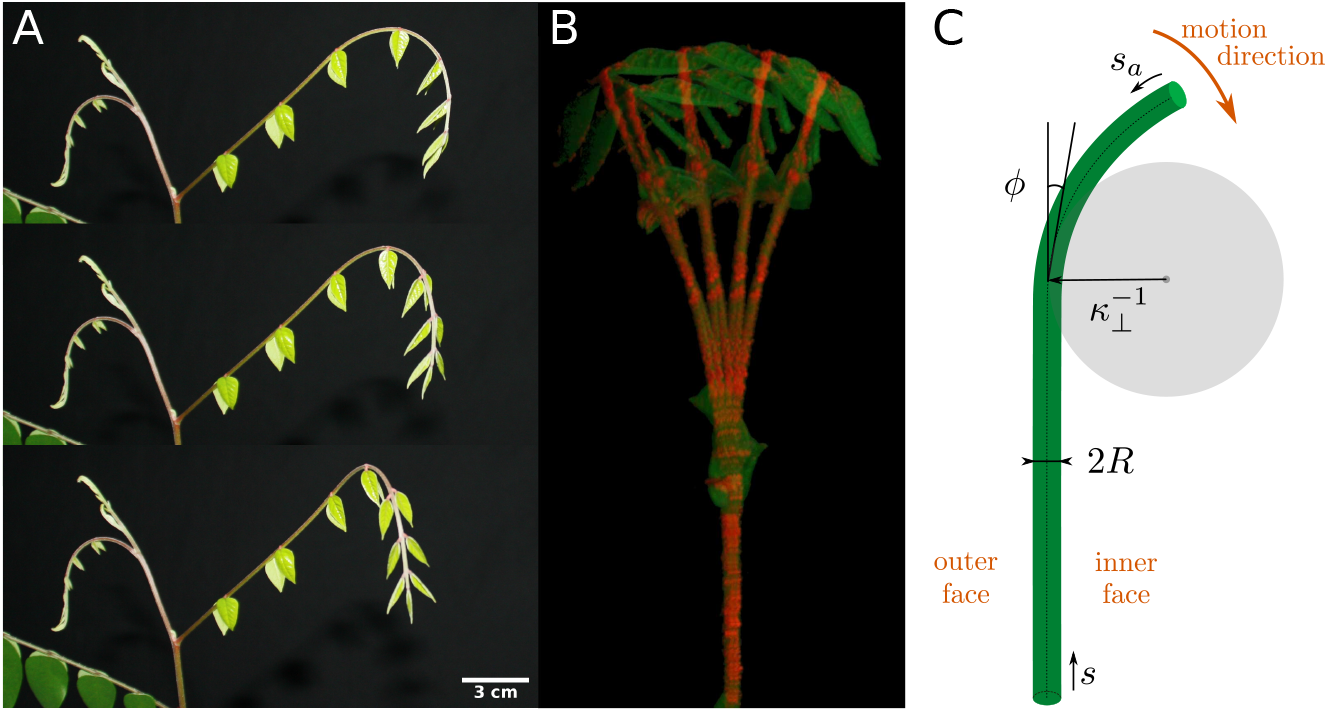
Time-lapse image of the nutation of a *Averrhoa carambola* leaf. **(A)** (side view). 30 minutes lapsed between each picture. One can see, from top to bottom, the hook coming out of the plane towards the observer **(B)** (top view). Eight superposed pictures taken every 15 min (the period of the motion is generally comprised between 1.5 and 4 hours). The leaf oscillates in a pendulum-like fashion, orthogonal to its growth axis, and superposing itself along the oscillation. At the end of one period the extension of the leaf is visible **(C)** Geometrical parameters describing the rachis and nutation.

### Elongation and bending are localized

We have evaluated the average elongation rate *Ė* of each of the successive interfoliolar segment by tracking the position of the successive nodes. The spatiotemporal diagram of *Ė* shows that only the apical-most region of the rachis elongates, defining a growth zone near the apex (see Fig. 2B). This is consistent with our previous findings on the same system (8). Here we also estimate the profile of differential elongation 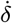 along the rachis from the transverse curvature measurement, thanks to the several hypotheses described in the Material and Methods section. We have then retrieved its envelope thanks to a Hilbert transform. The evolution in time and space of the envelope of 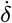 is displayed in Fig. 2C. We see that the differential growth—hence the bending—is spatially limited to a zone downstream of the apex. Similarly to what is done for the elongation, it is thus possible to define a bending zone. Our kinematics study reveals that this bending zone is at an approximately constant distance from the apex, similarly to the constant length of elongation zone from the apex (see Fig. 2C). Finally, going a step further in the description of nutation, we notice that the amplitude of the differential elongation—or of the bending—varies in time, reaching a maximum of 3 × 10^−2^ *h*^−1‡^.

**Fig. 2.**
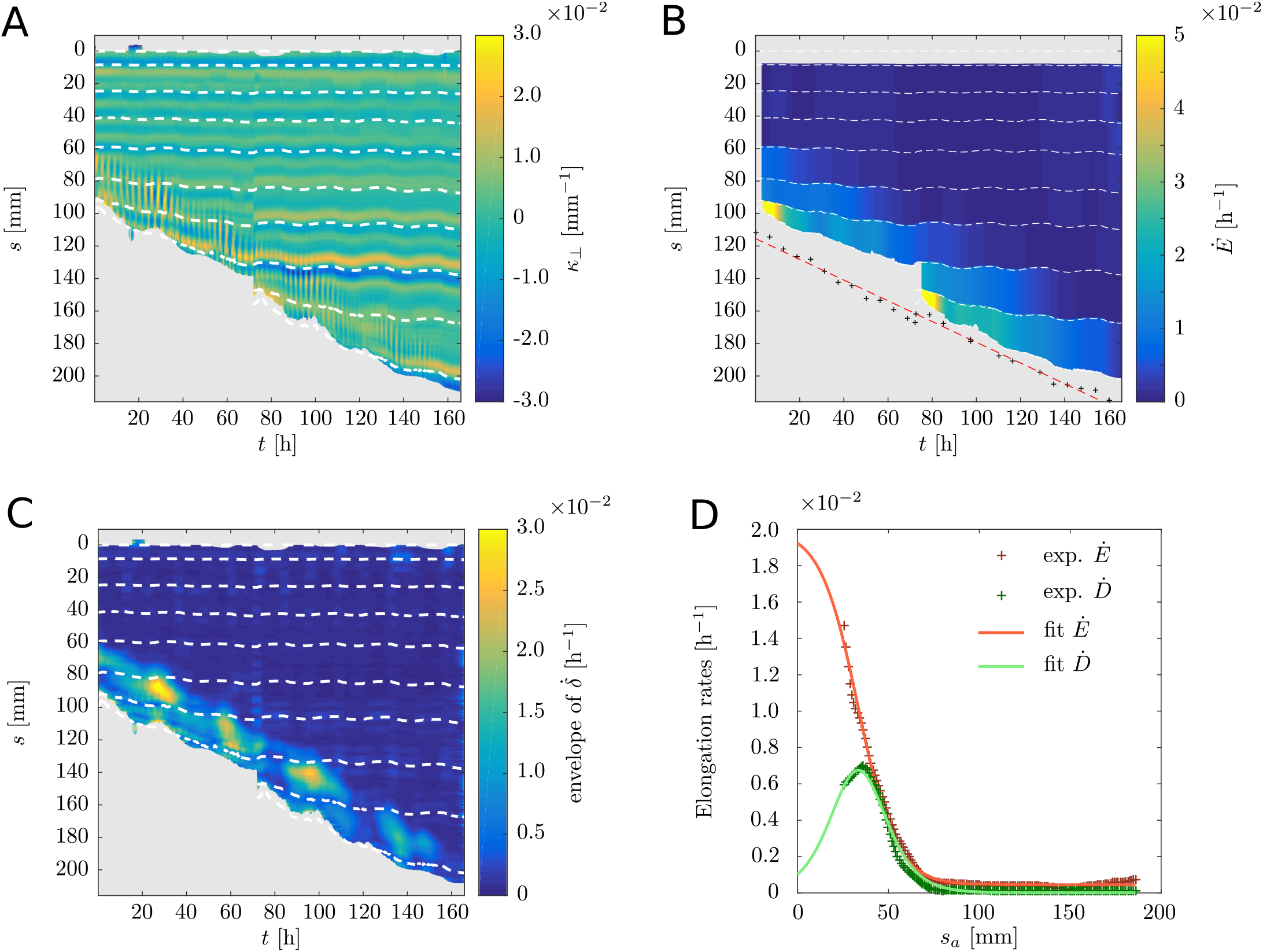
Growth kinematics underpinning the nutation motion in an *Averrhoa carambola* compound leaf. **(A)** Spatiotemporal diagram of the curvature *κ*_⊥_ along the rachis obtained from a top-view time lapse movie. The thin vertical yellow-blue stripes are the nutation oscillations. **(B)** Spatiotemporal diagram of the elongation rate *Ė* of each interfoliolar segment estimated from the leaflets’ trajectories (white dotted lines). The black crosses show the position of the leaf apex estimated from side-view pictures. The red dashed line is a linear fit of the apex position. **(C)** Spatiotemporal diagram of the envelope of differential elongation 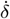 estimated from the curvature diagram (nutation amplitude).**(D)** Spatial profiles of elongation rate (red crosses) and differential elongation rate (green crosses) averaged over time. The two profiles were fitted respectively to a sigmoid (red line) and to its derivative (green line). The complete profiles cannot be measured from a top-view because of the leaf hooked shape.

### Differential elongation peaks where elongation drops

Because the growth spatial profile is almost steady in the frame of reference of the apex, we can average the measured quantities along the backward arc length *s*_*a*_. The averaged quantities *Ė* and 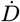 corresponding to mean elongation and differential elongation rates of interfoliolar segments are plotted on Fig. 2D. Both quantities confirm quantitatively the existence of a localized growth zone. The typical length scale is about 50 mm, and after 100 mm growth is not detectable at all. The mean elongation rate variation looks like a sigmoid function. In the growth zone the typical elongation rate is about 10^−2^ *h*^−1^, consistently to typical averaged values found in the literature (43, 44), and then drops to zero. Interestingly, the differential elongation rate behaves differently. It is nonmonotonous and its maximum coincides with the edge of the growing zone, where the mean elongation rate drops. A simple mathematical description of these sigmoid and peaked shapes is well fitted with the hyperbolic functions similar to Eq. 3 and 4. The results are displayed Fig. 2D. In this case the derivative of the fit of the longitudinal elongation rate matches well our experimental measurements of the differential elongation rate, with its amplitude remaining a free parameter (see supplementary text).

### The elongation profile in the growth zone is compatible with local contractions

We used techniques inspired from particle image velocimetry (see Materials and Methods) to quantify the elongation profile within the bending zone. The raw results shown in Fig. 3A only represent the *apparent* elongation rate 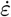 for the interfoliolar segment which contains the nutation zone. A nutation event is visible with the red-blue oscillation stripes. Because of the strong projection effects of the side view, the apparent elongation rate 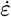 even turns negative (blue). The other striking feature is that, depending on *s*_*a*_, these oscillations have different periods: faster an the apical end (top on graph), and slower at the distal end (bottom on graph). This is confirmed quantitatively by the wavelet transform displayed in Fig 3B. We see distinct dominant modes with periods *τ*_2*f*_≈ 1.2 *h* and *τ*_*f*_≈ 2.1 *h* close to the apical (respectively distal) end of the studied rachis segment. To understand this observation, we developed a simple kinematics model. It is based on the measured experimental growth law: the data of Fig. 2D formalized by Eq. 3 and Eq. 4 (see Materials and Methods), and taking into account the effects of side view projection. This model also aims at understanding if the contractions seen in blue in Fig. 3A are only due to projection effects or correspond to real contractions on the side of the rachis. This model has at least two interesting outcomes. First it provides an order of magnitude for the differential growth. Indeed, we find that the parameters *R* (radius of the rachis), Δ*L* (length of transition of the sigmoid representing the growth zone),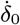 (amplitude of the differential growth), *ω* (pulsation of the nutation) and Δ*ϕ* (angular amplitude) are in fact constrained by the relationship

**Fig. 3.**
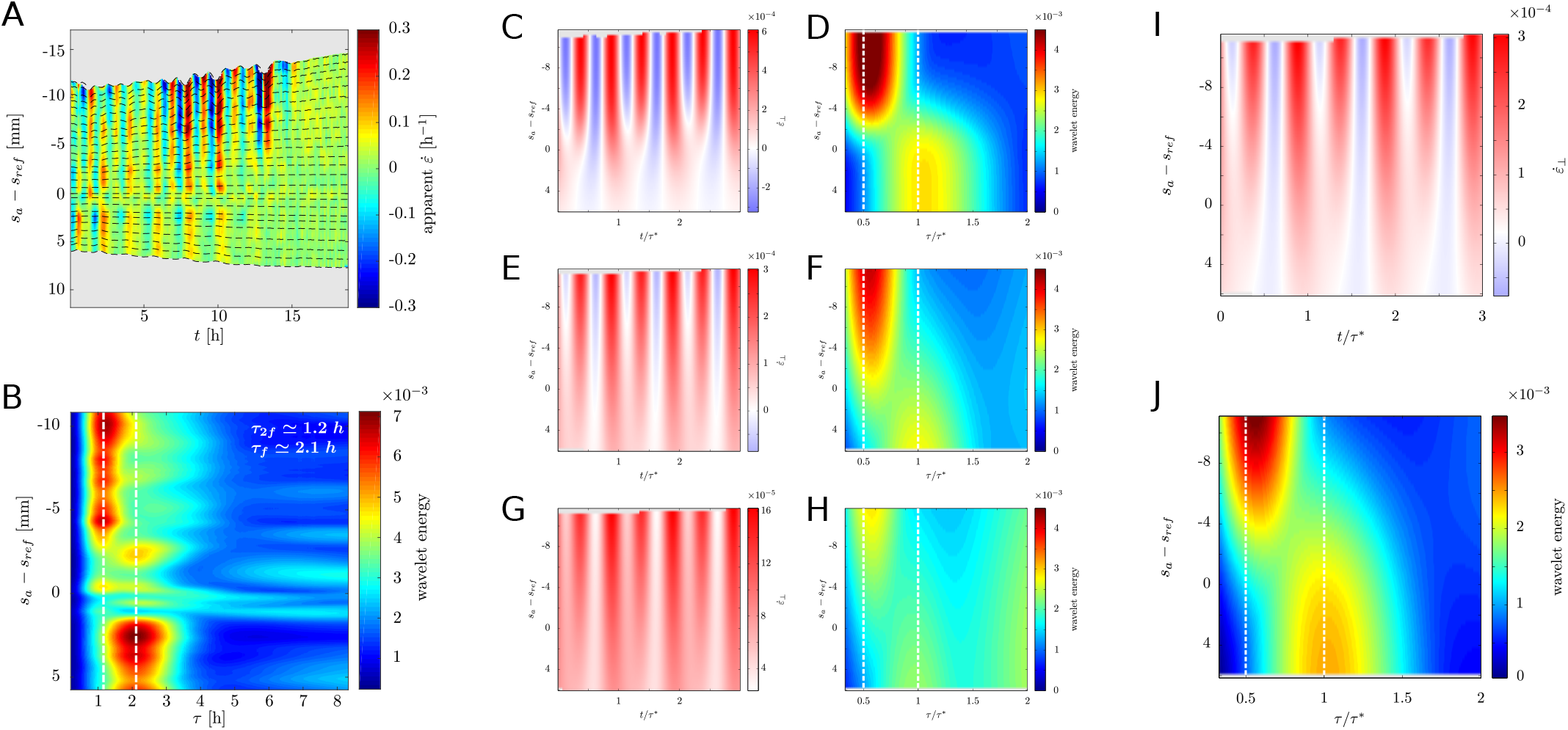
The comparison of our experimental data with our kinematics model suggests that nutation dynamics are compatible with local contractions. **(A)** Spatiotemporal diagram showing an experimental measurement of the *apparent* local elongation rate 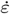 in the bending zone from a side-view time lapse movie. Because of the oscillatory motion of the rachis, the elongation rate measured is affected by projection effects. **(B)** Wavelet decomposition of the experimental spatiotemporal diagram of apparent elongation rate. The decomposition shows that two dominant modes in the signal: *τ*_2*f*_ ≈ 1.2 *h* and *τ*_*f*_ ≈ 2.1 *h* respectively close to the apical and basal ends of the observed section of the rachis. **(C-J)** Pairs of diagrams showing simulation outputs for different choices of parameters. The diagrams show apparent elongation rates (blue to red) and wavelet decomposition. **(C)** and **(D)**: 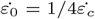 (possible contractions). **(E)** and **(F)**: 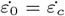 (threshold for contractions). **(G)** and **(H)**: 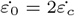 (no contractions). **(I)** and **(J)**: Best fit of the kinematics model to the experimental data; Δ*ϕ* = 8°, *L*_*gz*_ = 20.6 mm, Δ_*L*_ = 12.2mm, 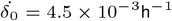(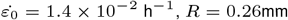, were fixed before fitting). This set of parameters allows local contractions.

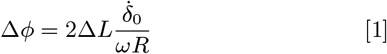

This can be understood as 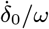 being the total differential growth over one period, which divided by the radius *R* gives the local curvature of the rachis, and integrated over the bending zone length 2Δ*L*, gives the final deviation of the apex. By plugging orders of magnitude in this relationship, one finds 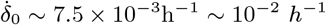^§^ which is the order of magnitude of the measured averaged growth, confirming the possibility of contractions. Second, by exploring the parameters space, we show that the observation of two distinct periods on the projection depends on the relative values of the differential elongation amplitude 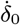 and the average elongation amplitude 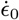 (see Fig. 3C-G). One can also fit the wavelet transform spatiotemporal diagram as a way to estimate the unknown experimental parameters. The best fit and the corresponding parameters are presented in Fig. 3C and D. They indicate that the rachis *must* locally contract to explain our experimental measurements.

### Asymmetric rigidity and cell wall composition in the bending zone

Building on our kinematics results and methods, we probed the microscopic properties of the rachis within the bending zone when the angle *ϕ* → 0, *i.e*. when 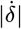 is expected to reach its maximum. We define the outer and inner faces of the rachis as backward and forward side of the nutation motion (see Fig. 1C). These faces are expected to have the largest, and respectively lowest, growth rates. The global mapping of a transverse cut of the rachis reveals an important variability of rigidity across the inner tissues (see Fig. 4A). The relative stiffness (normalized by the global average) across the rachis is widely distributed with a standard deviation of 52%.

**Fig. 4.**
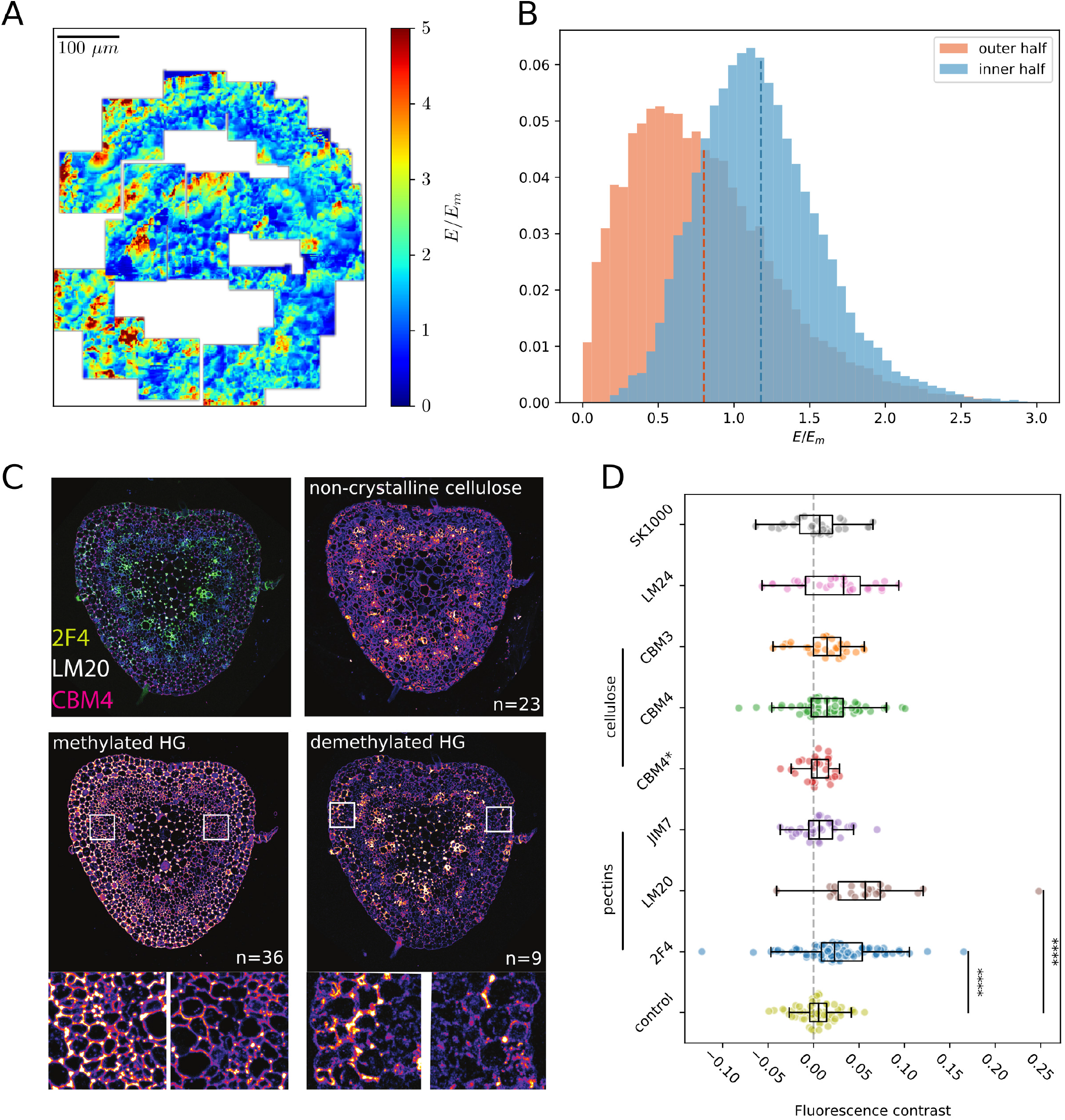
**(A)** Global rigidity mapping of a transverse cut of the rachis obtained by atomic force microscopy. **(B)** Relative rigidity histogram of the peripheral tissues in the bending zone. The histograms of the concave (blue) and convex faces of the rachis are drawn separately for comparison. Rigidity scores were normalized by the global mean on the sample. **(C)** Fluorescence images of transverse cuts from a single rachis in the nutation zone. Three histochemical treatments are presented together (composite image) and individually. The used antibodies target non-crystalline cellulose (CBM4) and homogalacturonans with a low/high methyl-esterification degree (2F4/LM20 respectively). For the HG images, two zoomed windows are presented for a detailed comparison of the lateral faces of the rachis. **(D)** Distributions of fluorescence contrast for several immunolabelling treatments. The contrast is calculated as (*< I*_*out*_ *>* − *< I*_*in*_ *>*)*/*(*< I*_*out*_ *>* + *< I*_*in*_ *>*), where *< I*_*X*_ *>* corresponds to the fluorescence intensity averaged over regions of interest located either on the inner or outer half of the rachis. Each dot represents a single measurement. The distributions are overlayed with their respective boxplot. Significant Student’s t-test have been indicated. The control measurement corresponds to the natural autofluorescence signal of *Averrhoa carambola*’s cell walls observed at *λ* = 405 *nm*.

To further check if the lateral faces of the rachis had different mechanical properties, we focused our mapping effort on the peripheral tissues—with thicker walls—where we would expect maximum differences. We quantified the distributions of relative stiffness of 3 different growing leaves in the nutation region (see Fig. 4B). Our results show that the tissues on the inner face of the rachis are on average 17% softer (standard deviation 6%) than the tissues on the outer face. Among the three first moments of the rigidity distributions, only the mean gave a consistent trend across biological repetitions (see Fig. S2 and Table S1 for repetitions and complete distribution characterizations).

Finally, we checked if the measured mechanical asymmetry correlates with biochemical asymmetry within the cell walls. We used multitarget immunolabelling to probe epitope asymmetry across the tissue (see Fig. 4C). We targeted several important families of components of the cell wall: cellulose, hemicelluloses and homogalacturonan domains of pectins. Additionally, the auto-fluorescence signal of *A. carambola* cell walls was used as a reference as it allows to check for any underlying anatomical asymmetry such as cell wall thickness for example. Results in Fig. 4D show the distributions of signal contrast for the different tested antibodies. Student’s t-test revealed no statistically significant bias in the auto-fluorescence signal (*p* = 0.13 *>* 5 × 10^−2^, see Fig. 4D). We then conducted a systematical comparison of antibody fluorescence and auto-fluorescence contrasts via Welch’s unequal variances t-test. Our results show no statistically significant shift from the reference signal for crystalline cellulose (CBM3), amorphous cellulose on pectolyase-treated samples (CBM4*) and mannans (SK1000). Xyloglucans (LM24) and regular amorphous cellulose treatments (CBM4) returned p-values of 1.3 ×10^−2^ and 2.8 × 10^−2^, advocating for a slight but unconclusive signal asymmetry. In contrast, both 2F4 and LM20 signals, marking HG with low and high degrees of methylesterification respectively, did significantly differ from the auto-fluorescence signal (both *p* < 5× 10^−5^ – for a complete description of the statistical tests results, please refer to Table S2). From the fluorescence contrasts, we see that the 2F4 and LM20 signals are on average 5% and 13% stronger on the outer half of the rachis than on its inner half.

## Discussion

### The nutation zone corresponds to the growing zone and undergoes “stop and go” phenomena

The kinematics data of the present study are consistent with our previous study indicating the presence of a steady growth zone, extending at a fixed distance from the apex (8). Here we show that the end of this growth zone also coincides with the nutation zone which is characterized by rapid changes of its curvature *K*_⊥_ due to large differential growth. Quantitatively, when and where the differential elongation rate is maximum, its amplitude is then comparable to the local mean elongation rate (see the obtained growth profile displayed Fig. 2D). This could be schematized as a “stop and go” phenomenon, where each side of the rachis grows alternately, before growth and motions cease altogether.

This coincidence of the maximum of the differential growth with the maximum of decrease of mean elongation is consistent with previous qualitative observations on *Arabidopsis thaliana* roots (45). This supports ideas of a separation between growth potential/signal and growth possibilities. At the apex, in the most growing zone, the growth potential (as influx of materials) or signal (as influx of hormones) is well above the growth possibilities (as maximum extension of the walls without breaking). Even if there would be variations in the potential/signal, in space or time, they would not be visible as the actual growth is already at its maximum, edging them out. On the contrary when the growth potential or signal globally starts to decline, near the end of the growth zone, then any variation in the growth potential or signal is directly translated into a variation of actual growth. The same interpretation could apply to the straightening of wheat coleoptiles (46): as the coleoptile bends towards the vertical, the differential growth potential or signal is at its maximum, and no oscillation is observed. On the contrary, when the coleoptile approaches a vertical posture, the potential or signal is lower, and nutation of the tip becomes visible again.

### Correlation between growth and cell wall properties

Our results belong to a long series of observations correlating growth with changes in cell wall elasticity, by suggesting that growth is enhanced where the Young’s moduli are lower. This phenomenon was evidenced in growing pollen tips (47), maize roots elongation zone (48, 49), *Arabidopsis* shoot meristem before primordia formation (50, 51) as well as *Arabidopsis* hypocotyls growth where anisotropic growth was associated to longitudinal walls softer than transverse ones (52). In this last study, as well as in a few others (51, 53–55), the change in elastic properties was attributed to changes in chemical state of the pectins. The de-methylesterification of the pectins would trigger elastic softening (51, 56, 57), consistently with the idea that the de-methylesterified form is a target for pectin-degrading enzymes, such as polygalacturonases, affecting the texture and rigidity of the cell wall (58).

Therefore, the changes in cell wall elasticity associated with growth could only be an epiphenomenon of the change in density of the cell wall associated to the expansion of the homogalacturonans (HG) following their de-methylesterification. Note that the orders of magnitude of relative changes are the same for pectin content and Young modulus (of the order of ten percent in both cases). Fully grasping the link between HG chemistry and growth is tempered by the dynamics of its metabolism as discussed in (59). The immunolabelling approach can only observe one frozen state. Thanks to the oscillatory nature of the growth studied in the present study, we could capture and compare two events simultaneously: contraction and expansion of the tissue. The shift from one condition to the other would typically happen in 30 minutes. Interestingly, the most prominent difference we observed in cell wall composition across the tissues is in the highly methylated and the highly demethylated HG. These two components were recently demonstrated to form nanofilaments controlling the anisotropic elongation linked to pavement cell morphogenesis (38). Pectins have multiple ways of action (34). They could also form Ca^2+^ bonds, which promote the formation of the so-called ‘egg-box’ model structure, thus forming gels (60) and increasing elasticity. This was observed for tip-growing pollen tubes where a lower degree of methylation of HG was attributed to the stiffer cell wall parts, while higher degrees of methylation was observed in softer parts (61, 62). This is not the case here, as the partially demethylated HG epitopes detected by JIM7, which present a gel-like structure, do not seem to be implicated in growth processes (38, 63).

### The hidden link between microscopic properties and macroscopic effects

In our system it is difficult to disentangle the reversible and irreversible contributions to growth as it was done by Proseus *et al*. for the single-cell algae *Chara* (64). It has also been shown in the case of the shoot apical meristem that elastic inhomogeneities (or differences in stress stiffening) could lead to differential growth (40). Therefore, to discuss the missing link between the observed microscopic properties and the macroscopic contractions, we propose two different hypotheses.

First, one should consider the reversible processes as they have already been found to be involved in nutation. Local contractions have indeed been measured at the organ level (65– 67) and at the cellular level (68, 69). This last paper actually shows that individual cells in the bending zone of *Phaseolus vulgaris* undergo reversible volume variations; this clearly advocates for water movements. Recently, Cheddadi *et al*. formalized the water fluxes coupling in multicellular organs and showed that new types of lateral inhibitory mechanisms could amplify growth heterogeneities (70). The quantitative analysis of Cheddadi’s model in the case of the geometry of a bilayer goes beyond the scope of this paper but numerical simulations indicate that elasticity variation cannot be enough to induce oscillations alone. The main explanation of this being that elasticity hardly gives the relaxation timescale of pressure. Overall, direct observations of water movement events are scarce and the mechanism that could control them would require symplasmic and periplasmic water movement control and are still elusive.

From our observation, one can propose a second hypothesis for the temporal events: on the growing side, HG are actively addressed to the cell wall in their native methylated way. Then growth turns to the other side of the rachis and HG are sparsely degraded/recycled by endoglucanase explaining the reduction in staining observed in the methylated and non-methylated pectins. Such recycling was proposed for other tissues. Here we can indicate that the time scale could be as fast as 30 minutes. As discussed in (38), the expansion part could be solely due to HG filament expansion following the de-mehtylesterification. In addition the partial removal of the highly charged polymer following their recycling could as well lead to cell wall compaction in link with the observed tissue contraction.

## Conclusion

In conclusion, we have described quantitatively the phenomenon of nutation in *Averrhoa carambola* with data collected at several scales. At the scale of the organ, we have shown that nutation coincides with the growing zone involved in some posture regulation processes (8). We deduced from kinematics a *growth law* which shows that differential growth happens at the edge of the growing zone where mean elongation rate collapses. By focusing on this zone, our particle image velocimetry technique reveals a “stop and go” phenomenon for growth implying even local contraction of the non growing region. Finally the microscopic investigation revealed that the key difference between the convex and the concave sides is the cell wall biochemistry of the HG. Yet the link between these microscopic finding and the possibility of contraction is still an open question but our observation opens new perspective for the understanding of growth processes in plants.

## Materials and Methods

### Growth conditions of *Averrhoa carambola*

The conditions of culture are the same than in (8). *Averrhoa carambola* seeds were directly obtained from commercially available fruits and sown into all-purpose compost. Young seedlings were first kept inside a small lab greenhouse. Older plants (*>* 6 months) were then moved to the experimentation room. There, plants were submitted to a 12/12 light cycle under ORTICA 200W 2700K culture lamps. The temperature and relative humidity rate were monitored with a DHT22 sensor. Temperature was usually comprised between 20 °*C* and 24 °*C*. The relative humidity rate was around 60%.

### Kinematics: sample preparation

The rachis of interest was carefully coated with fluorescent pigments with a brush. For curvature and coarse elongation measurements, the top of the rachis was coated homogeneously with orange pigments. Small blue fluorescent dots were added to mark the nodes and the petiole. For fine measurements of local growth, the orange pigments were deposited on the face of a few interfoliolar segments so that they form highly textured and contrasted patterns. In both cases, because of growth, pigments needed to be added manually on a regular basis to compensate the dilution of the signal over time.

### Kinematics: image acquisition

The kinematics of nutation were captured using time-lapse photography with a DSLR camera controlled with the open-source software gPhoto2. The camera was firmly fixed to a rigid structure to avoid any displacement or rotation. The built-in flash of the camera was covered with LEE Moss green filter and set to the lowest intensity to keep light input minimal during nights. For curvature and coarse growth kinematics, top-views were taken every 2.5 min. For local growth measurements, side-views were taken every minute.

### Kinematics: data analysis

The midline (or skeleton) of the rachis was obtained by first thresholding the red channel of the pictures. A cloud of point was then obtained and reduced to a smooth line with a moving median filter. The curvature of the rachis *κ*_⊥_ in the plane of interest was obtained by locally fitting the midline to a circle. The position of the leaflets was retrieved by thresholding the blue channel. Because of growth, blues dots dilated, lost intensity in time and sometimes even split. The global unfurling motion of the rachis sometimes resulted in a temporary occlusion of some blue dots. Simple rules on the conservation of these dots, distance between consecutive dots and displacements values could overcome a majority of tracking failures. Manual correction was still needed in some special cases. Finally, the presented spatiotemporal graphs were smoothed with 2D averaging and median filters.

### Kinematics: fine elongation measurements

We obtain the skeleton of the rachis by a simple geometric transformation of the upper contour which is less altered by leaflet motions. Then we measure the the elongation field along the rachis by using image-to-image correlation algorithm similar to (71). The time-frequency analysis of the elongation signals was done by using MATLAB’s continuous wavelet transform toolbox. We used the ‘cgau2’ mother wavelet (second order derivative of the complex Gaussian). For each location of the rachis, 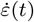 was wavelet-transformed. From the resulting complex coefficients *C*_*a,b*_ we extracted information on the weight of each scale/frequency in the signal by computing an ‘energy’: *E*(*a*) =∑ _*b*_ | *C*_*a,b*_|^2^ / ∑_*a*_ ∑_*b*_ | *C*_*a,b*_ |^2^, where *a* and *b* are the scale and shift parameters of the wavelet transform. This information was then re-aggregated and re-arranged to build kymographs displaying the weight of frequencies in the elongation signal along the rachis.

### Details and implementation of the model

The rachis is modelized by a two-dimensional beam of width 2*R* (see Fig. S3) and of total length *L*_*tot*_. The geometry of the midline is then described with the same quantities than the actual leaf (see Fig. 1C). The model contains only a few essential ingredients:

1. The lateral faces of the beam can have different elongation rates 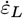 and 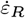, giving rise to differential elongation 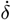. We assume that the profile of elongation is linear in the bulk of the rachis:

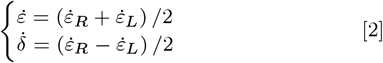
2. An apical growth zone of length *L*_*gz*_. The elongation rate of the midline 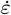 as a function of the reverse arc length *s*_*a*_ at time *t* is thus given by:

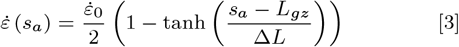
3. Differential elongation occurs where the mean elongation rate drops, within a bending zone of length 2Δ*L*. Because nutation is a periodical oscillatory motion, differential elongation is modulated by a sinus of frequency 2*π/ω*:

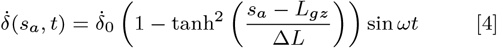
4. We assume differential elongation is the unique driver of the bending of the rachis. In our case, since the period of nutation is much smaller than the typical time scale of elongation, we furthermore neglect the advection of curvature and write:

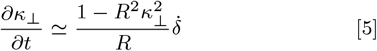

The model was implemented numerically with discretized versions of the kinematic Eq. (3), Eq. (4) and Eq. (5).

### Atomic force microscopy assays

Studied leaves were filmed overnight before the AFM experiment in the same conditions than for our kinematics experiments. Leaves were cut when *ϕ* ≃ 0°, when the difference in the elongation rates of the lateral faces is expected to reach a maxima. Several samples were taken free-handedly from the nutation zone with a razor blade. We kept track of the orientation of samples by resorting to several anatomical clues. The distal/apical axis was checked thanks to fact that trichomes consistently point toward the apex. The adaxial/abaxial axis was checked thanks to the bilateral symmetry of the rachis (see Fig. S4). Samples were then placed vertically on a microscopy slide and partially embedded in agarose like described in (57). Samples were then kept immersed for the whole experiment in a solution of mannitol for plasmolysis. Measurements began after 20–30 min so that plasmolysis was reached (see Fig. S5). Identation was achieved with cantilevers with a spherical bead of diameter 25 *µm*. The indentation depth is not directly controllable but was generally comprised between 1*µm* and 5*µm*. Force-distance curves were analyzed with the proprietary JPK software to extract Young moduli with Hertz contact model. We chose to work with relative Young moduli (normalized by the average Young moduli of the sample). The AFM mapping of the the sample was achieved by repeatedly indentating over a 100 *µm ×* 100 *µm* probing zone, which was manually moved to cover the regions of interest. Some aberrant points (impurities, trichomes) were withdrawn thanks to simple filters on relative height and stiffness values. Measurements outside falling outside the surface of the rachis surface were manually masked.

### Immunolabelling experiments

Samples were taken as described in the previous section. Then, we performed multitarget immunohistochemistry following a recently published protocol (72). First, samples are gradually dehydrated and biologically fixed in a bath of paraformaldehyde. Then samples were embedded in wax. Samples were sliced (thickness = 5 *µm*) with a microtome, mounted on microscopy slides, dewaxed and progressively rehydrated. Samples were then placed in a buffer which nature depends on the chosen antibodies. The primary antibody is injected over the microscopy slide and let to react overnight. After washing the samples with the adequate buffer, the secondary antibody is introduced and let to react for at least 12 *h* in darkness, to avoid bleaching. Finally, the samples are washed a last time and the microscopy slides sealed. Observations were made under a confocal microscope. The following antibodies were used: 2F4, JIM7, LM20, CBM3, CBM4, LM24 and SK1000. These bind respectively: pectins with low degree of methylesterification (DM), high DM pectins, high DM pectins, crystalline cellulose, amorphous cellulose, xyloglucans and mannans. The additional CBM4* treatment uses CBM4 antibodies after treating the samples with pectolyases to free the cellulose epitopes. Finally, we used the natural autofluoresnce signal of Averrhoa carambola’s cell walls observed at *λ* = 405 *nm* as a reference signal. Raw confocal images were processed with the ImageJ software. Each z-stack was reduced to a simple multichannel image with a maximum intensity projection. The transmission channel was used to discriminate the tissues of interest. For each sample, the peripheral tissues (excluding the central pith, vessels and outer epidermis) were divided in two arc-shaped regions according to the bilateral symmetry of the rachis. The intensity of the multiple fluorescence signals were averaged over these regions of interest. Finally, a constrast score was computed for each signal and for each sample as: (⟨*I*_*out*_ ⟩ − ⟨*I*_*in*_⟩) */* (⟨*I*_*out*_⟩ + ⟨*I*_*in*_⟩), where *I*_*X*_ stands for the fluorescence intensity of either side.

## Data Availability

All study data are included in the article and supporting information.

## ACKNOWLEDGMENTS

The authors thank Elliot Meyerowitz and Raymond Wightman for their support and help on the immuno-labelling experiments. We are also grateful to Olivier Hammant and Emmanuel de Langre for continuous feedback on our works. MR is grateful to “Ecole Doctorale Frontières du Vivant — Programme Bettencourt” for financial support.

## Footnotes

* The definition slightly evolved in time. Bonnet discussed nutation in the context of phototropic movements. Sachs redefined this term later in a more general context mentioning “curvatures caused by the unequal growth of different sides of an organ” (16)

† An alternative arc length *s*_*a*_ going the opposite way will also be used.

‡ However, these slow modulations of nutation are not in the scope of the present study

§ We used ΔΦ ∼ *π/*6, 2*π/ω* ∼ 2h, *R* ∼ 0.25mm and Δ*L* ∼ 50mm for all the presented simulations

