## Supplemental information for "Spatiotemporal growth pattern during plant nutation implies fast dynamics for cell wall mechanics and chemistry: a multiscale study in *Averrhoa carambola*"

#### This PDF file includes:

Supplementary text
Figs. S1 to S5
Tables S1 to S2
Legend for Movie S1

#### Other supplementary materials for this manuscript include the following:

Movie S1

### Supporting Information Text

**Details on the fitting procedure for $\dot{E}$ and $\dot{D}$.** The coarse elongation rate $\dot{E}$ and the coarse differential elongation rate $\dot{D}$ discussed in the Results section, and shown in Fig. 2D have been fitted to the following functions:

$$\dot{E}(s) = \frac{\dot{E}_0}{2}\left(1 - \tanh\left(\frac{s_a - L_{gz}}{\Delta L}\right)\right) \quad [1]$$

$$\dot{D}(s) = \dot{D}_0\left(1 - \tanh^2\left(\frac{s_a - L_{gz}}{\Delta L}\right)\right) \quad [2]$$

where $s_a$ is the arc length defined from the apex, $L_{gz}$ is the length of the growth zone and $\Delta L$ is the typical length scale of variation of the elongation rate. The $\dot{D}$ function here is proportional to $\partial_s \dot{E}$. The two functions were fitted together to the experimental data, in a single process with shared parameters $L_{gz}$ and $\Delta L$. The amplitudes of the functions were left independent of each other.

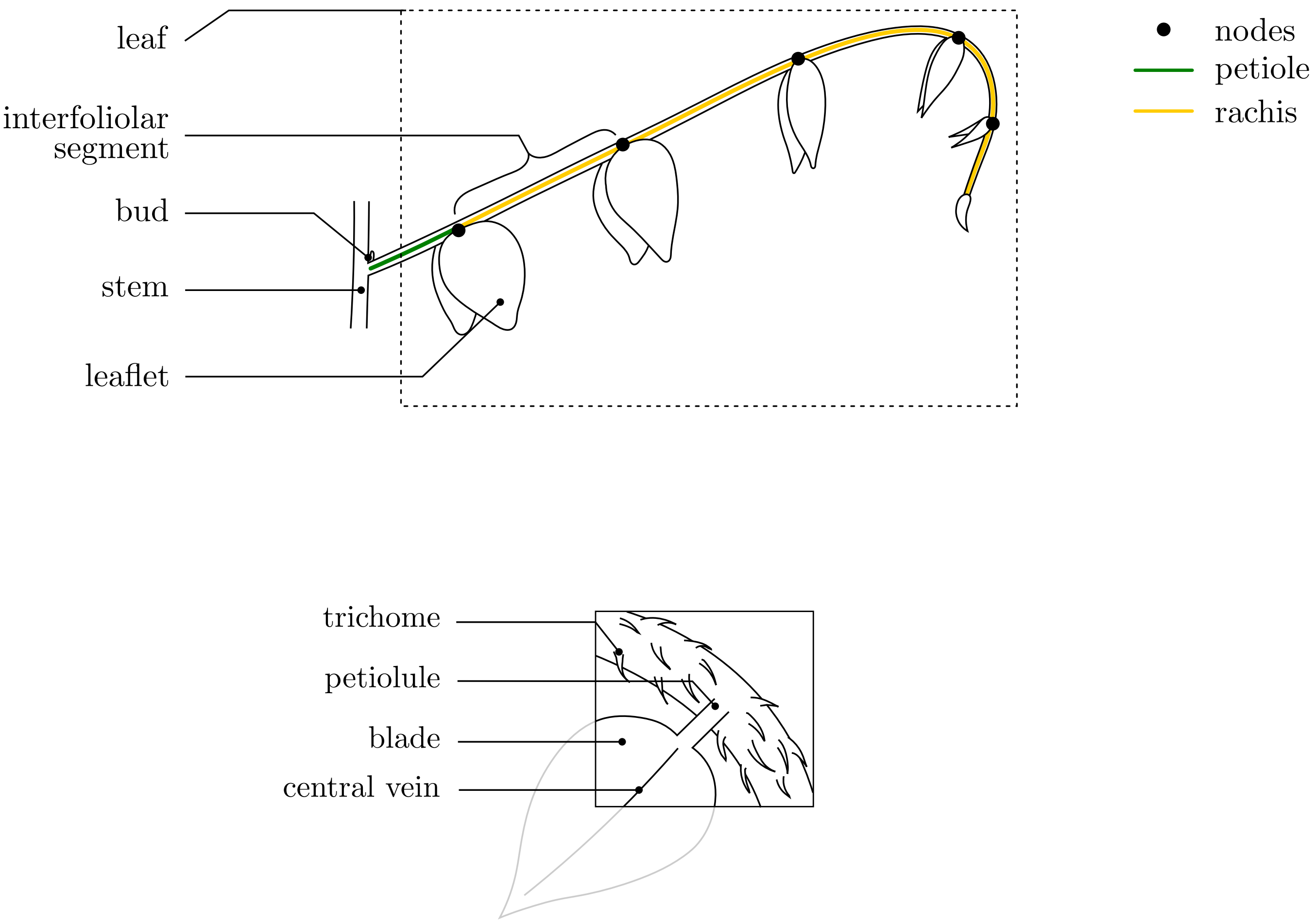


**Fig. S1.** Drawing of a growing *Averrhoa carambola compound* leaf. The top panel represents the entirety of a compound leaf and details the associated vocabulary. The bottom panel is a close-up around a leaflet, with additional details and anatomical vocabulary. The unit of interest for this study is the rachis, which can be seen as the “center vein" of the compound leaf ; and its subunits: the interfoliolar segments, separated by nodes where leaflets are connected to the rachis.

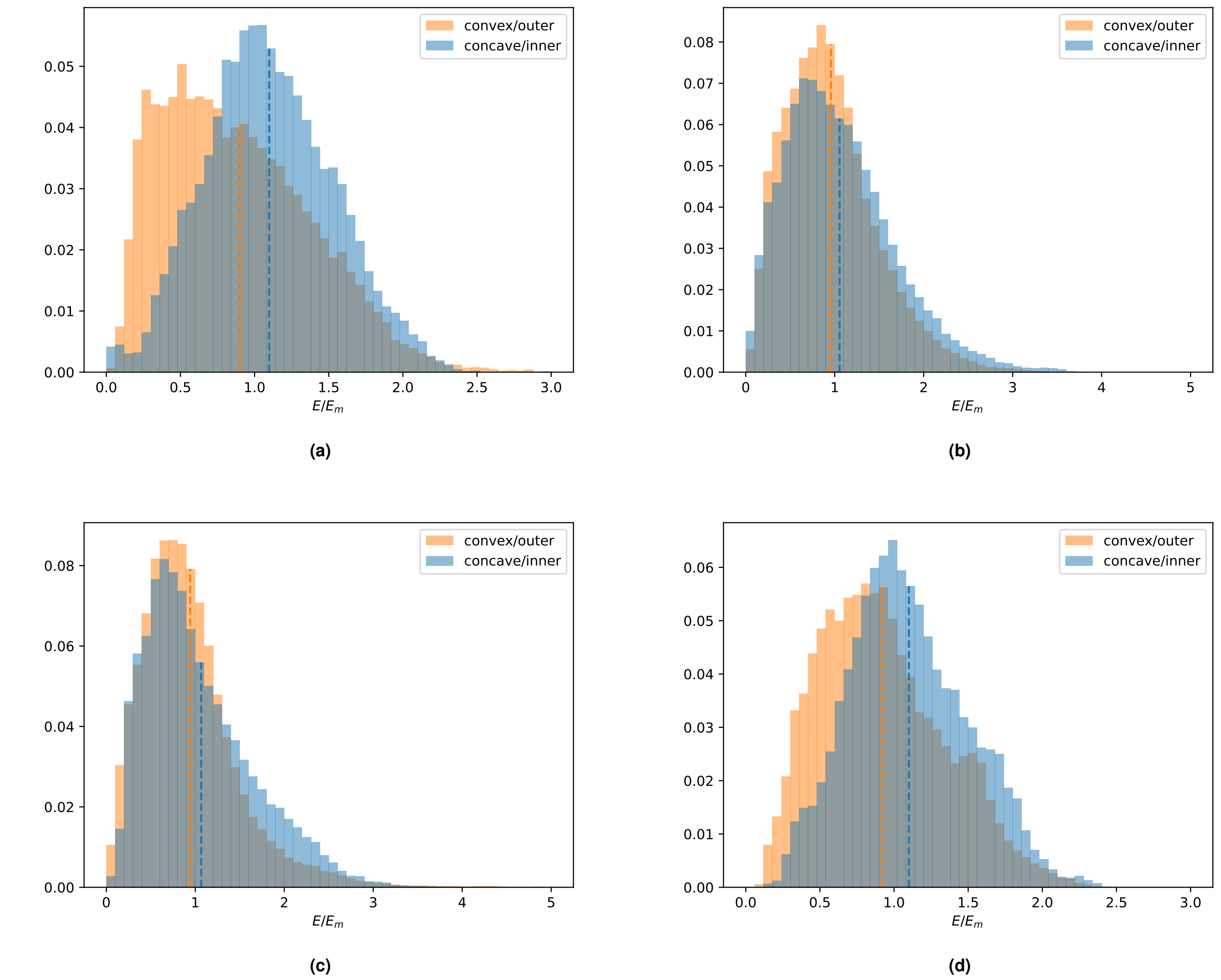


**Fig. S2.** Distributions of relative stiffnesses for our repetition experiments. Histograms of the convex/outer (orange) and concave/inner (blue) faces of the rachis are drawn separately for comparison. The dotted lines correspond to the means. (a) Distribution for another sample from the same rachis than data presented in Fig. 4B. (b-c) Two samples from a second biological repetition. (d) A single sample from a third experiment.

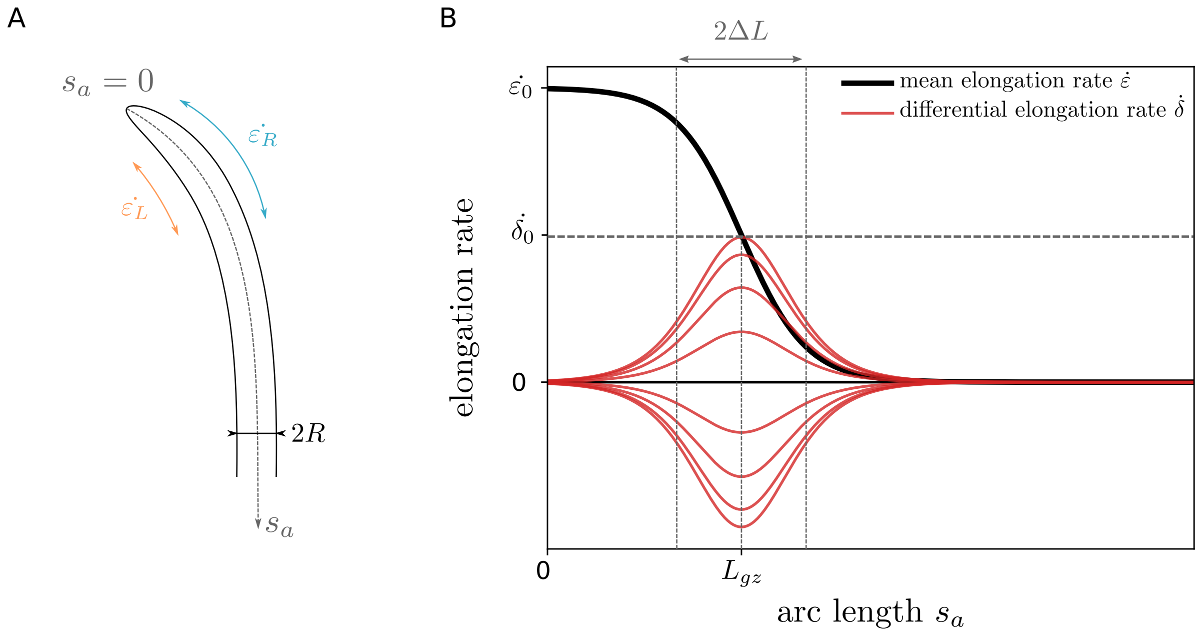


**Fig. S3.** (A) Geometrical parametrization of the model. (B) Elongation and differential elongation laws. The elongation rate $\dot{\varepsilon}$ shows a growth zone of length $L_{gz}$ defined from the apex. The differential elongation rate $\dot{\delta}$ takes place where elongation is dropping. It is proportional to the derivative of $\dot{\varepsilon}$. The differential elongation is furthermore modulated in time by a sine function of angular frequency $\omega$. Both functions are defined with hyperbolic functions, as discussed in the Methods section. Both quantities have a characteristic length of variation $\Delta L$

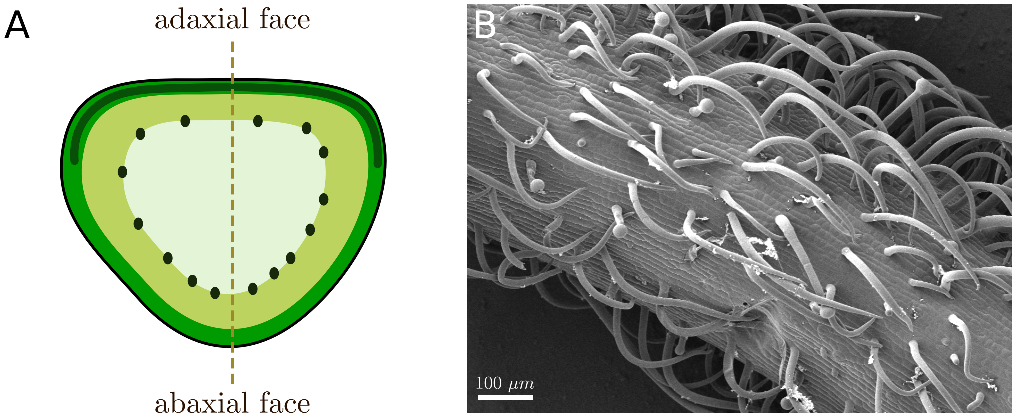


**Fig. S4.** Useful clues to determine the orientation of a sample. (A) Simplified picture of a cross-section of *Averrhoa carambola* rachis. The bilateral symmetry of the rachis allows a clear distinction between the lateral faces and the abaxial/adaxial faces. Flatness, chloroplast and vessel densities allow to distinguish the adaxial and abaxial faces from one another. (B) Cryo-SEM image of a carambola rachis. The rachis is covered with trichomes that all point toward the apex

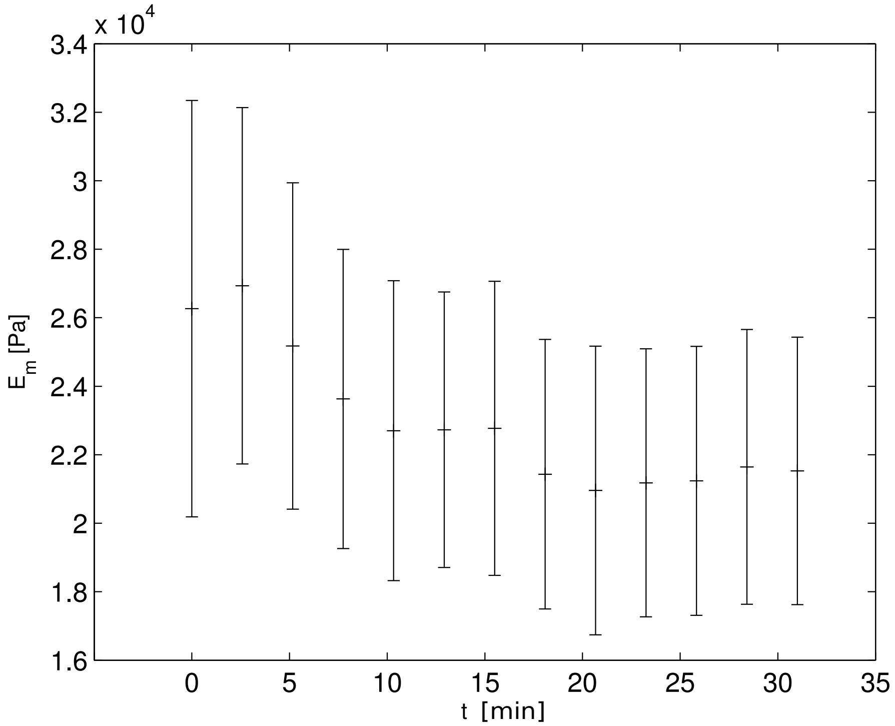


**Fig. S5.** Plasmolysis dynamics and plasmolysis effects on Atomic Force Mi- croscopy measurements. The same region is scanned repeatedly and the evolution of its average Young's modulus Em is traced. Error bars represent the standard deviation over the AFM map.

**Table S1. Summary of the quantification of the three first moments of the relative stiffness distributions.**

| Figure | Experiment | Sample | $\mu_{in}$ | $\mu_{out}$ | $1-\mu_{out}/\mu_{in}$ | $\sigma_{in}$ | $\sigma_{out}$ | $1-\sigma_{out}/\sigma_{in}$ | $\gamma_{in}$ | $\gamma_{out}$ | $1-\gamma_{out}/\gamma_{in}$ |
|---|---|---|---|---|---|---|---|---|---|---|---|
| Fig. 4B | 1 | 1 | 1.10 | 0.90 | $1.80 \cdot 10^{-1}$ | 0.43 | 0.56 | $-2.87 \cdot 10^{-1}$ | 0.16 | 1.77 | $-9.99 \cdot 10^{0}$ |
| Fig. S2a | | 2 | 1.18 | 0.80 | $3.20 \cdot 10^{-1}$ | 0.42 | 0.49 | $-1.75 \cdot 10^{-1}$ | 0.75 | 0.91 | $-2.01 \cdot 10^{-1}$ |
| Fig. S2b | 2 | 1 | 1.05 | 0.96 | $9.26 \cdot 10^{-2}$ | 0.61 | 0.54 | $1.13 \cdot 10^{-1}$ | 0.87 | 1.93 | $-1.21 \cdot 10^{0}$ |
| Fig. S2c | | 2 | 1.07 | 0.94 | $1.16 \cdot 10^{-1}$ | 0.87 | 0.71 | $1.83 \cdot 10^{-1}$ | 12.95 | 10.28 | $2.06 \cdot 10^{-1}$ |
| Fig. S2d | 3 | 1 | 1.10 | 0.92 | $1.64 \cdot 10^{-1}$ | 0.40 | 0.42 | $-5.72 \cdot 10^{-2}$ | 0.32 | 0.50 | $-5.46 \cdot 10^{-1}$ |

Subscripts *in* and *out* correspond to the inner (or concave) and outer (or convex) faces of the rachis, respectively. Remember that our working hypothesis is that, as the rachis bends, its outer face grows faster than the outer one. Individual experiments correspond to a single rachis. Samples correspond to different transverse cuts of a given rachis.

**Table S2. Summary of the statistical tests applied to immunolabelling experiments.**

| | Average contrast | std contrast | Student's t-test t-value | p-value | Welch's t-test t-value | p-value | $N$ |
|---|---|---|---|---|---|---|---|
| 2F4 | $2.78 \cdot 10^{-2}$ | $4.22 \cdot 10^{-2}$ | $6.17$ | $2 \cdot 10^{-8}$ | $-4.55$ | $1.18 \cdot 10^{-5}$ | $89$ |
| CBM3 | $1.29 \cdot 10^{-2}$ | $2.51 \cdot 10^{-2}$ | $2.8$ | $8.75 \cdot 10^{-3}$ | $-1.67$ | $0.1$ | $31$ |
| CBM4 | $1.58 \cdot 10^{-2}$ | $3.46 \cdot 10^{-2}$ | $3.46$ | $1.04 \cdot 10^{-3}$ | $-2.23$ | $2.79 \cdot 10^{-2}$ | $58$ |
| CBM4$^{*}$ | $4.04 \cdot 10^{-3}$ | $1.56 \cdot 10^{-2}$ | $1.35$ | $0.19$ | $-3.33 \cdot 10^{-3}$ | $1$ | $28$ |
| JIM7 | $7.18 \cdot 10^{-3}$ | $2.30 \cdot 10^{-2}$ | $1.74$ | $9.18 \cdot 10^{-2}$ | $-0.64$ | $0.52$ | $32$ |
| LM20 | $6.00 \cdot 10^{-2}$ | $5.03 \cdot 10^{-2}$ | $5.84$ | $5.04 \cdot 10^{-6}$ | $-5.28$ | $1.41 \cdot 10^{-5}$ | $25$ |
| LM24 | $2.37 \cdot 10^{-2}$ | $3.91 \cdot 10^{-2}$ | $3.32$ | $2.35 \cdot 10^{-3}$ | $-2.59$ | $1.35 \cdot 10^{-2}$ | $31$ |
| SK1000 | $4.73 \cdot 10^{-3}$ | $3.12 \cdot 10^{-2}$ | $0.79$ | $0.44$ | $-0.11$ | $0.91$ | $28$ |
| control | $4.02 \cdot 10^{-3}$ | $2.05 \cdot 10^{-2}$ | $1.52$ | $0.13$ | | | $61$ |

Here, $N$ corresponds to the total number of different samples tested. These samples are taken from 2 independant biological repetitions, on the interfoliolar segment where nutation occurs and the the two adjacent ones.

**Movie S1.** **Fourty-eight hours of development of an *Averrhoa carambola* compound leaf shown through synchronized top and side views. Nutation, the swinging motion of the rachis, is most easily observed in the top-view. The side-view offers a complete vision of growth motions of a compound leaf, with the typical hook shape and unfurling motion. Scale bars = 1 cm.**
